# Diversity, Equity, and Inclusion in the Melanoma Research Community

**DOI:** 10.1101/2022.09.24.509278

**Authors:** Marie E. Portuallo, David Y. Lu, Gretchen M. Alicea, Joel Bolling, Rebecca Lee, Jennifer McQuade, Allison Betof Warner, Michael Davies, Ashani Weeraratna, Jessie Villanueva, Vito W. Rebecca

## Abstract

Last October, within the 2021 SMR congress, we held the inaugural Diversity in Science Session. The goal of the session was to discuss diversity, equity, and inclusion in the melanoma research community and strategies to promote the advancement of underrepresented melanoma researchers. An international survey was conducted to assess the diversity, equity, and inclusion (DEI) climate among researchers and clinicians within the Society for Melanoma Research (SMR). The findings suggest there are feelings and experiences of inequity, bias, and harassment within the melanoma community that correlate with one’s gender, ethnic/racial group, and/or geographic location. Notably, significant reports of inequity in opportunity, discrimination, and sexual harassment demonstrate there is much work remaining to ensure all scientists in our community experience an academic workplace culture built on mutual respect, fair access, inclusion, and equitable opportunity.

## Introduction

> “Not everything that is faced can be changed, but nothing can be changed until it is faced”- James Baldwin

Structural oppression, an inequitable atmosphere, biased practices, and lack of inclusivity have historically plagued arenas of higher education and the academic community, with plentiful evidence that these issues continue to persist. To better understand the experiences with and perceptions of diversity, equity, and inclusion (DEI) within the melanoma researcher and clinician community, the Society for Melanoma Research (SMR) performed a diversity, equity, and inclusion (DEI) climate survey in the Summer of 2021. Results from this survey will provide insights into ongoing DEI challenges faced by members the SMR community and help to focus future DEI efforts to truly foster a diverse, equitable, and inclusive SMR community.

## Methods

### Participants

The Society for Melanoma Research (SMR) climate survey was distributed to more than 5,000 members globally to the SMR community, receiving 118 responses. Participants were asked about their gender, ethnic/racial group, sexual orientation, and continent of current residence with the option to answer these questions using their own terminology. Participants largely responded as White, Black, Asian, or Hispanic for their ethnic/racial group, male or female for their gender, and heterosexual or homosexual for their sexual orientation. The continent of participants’ current residence was divided into United States, Europe, Australia, and Other (answers from South America, Africa and Asia were pooled due to low number of respondents to allow for statistical analyses) after all responses were collected. Where denoted, N refers to the total number of participants from that particular group that responded to that question.

### Statistical Methods

Frequency tables were created to summarize survey responses for each question, and Fisher’s exact test was performed to determine nonrandom associations between groups and responses.

## Results

### Inequity in Opportunities

Respondents were asked to attest whether they felt inequity for opportunities in their local lab/clinic or at the institutional/university level. In regard to ethnic/racial group, 44.8% of non-white respondents (N=29) felt they missed out on opportunities in their lab/clinic due to their race/ethnicity relative to among only 8.4% of their white counterparts (N=83) (P value < 0.001) (**Table 1**). Additionally, 39.3% of non-white respondents (N=28) reported feeling a lack of opportunity in the lab/clinic due to their gender, regardless of gender; whereas only 18.1% of their white counterparts (N=83) reported the same feeling (P value = 0.037).

These reports of inequity were also consistent at the departmental/institutional level. 34.5% of non-white respondents (N=29) reported feeling a lack of opportunity in the department/institution due to their race/ethnicity versus 12.2% of white respondents (N=82) (P value = 0.011) (**Table 1**). In regard to geographical region, 34.3% of North American (N=70) versus 66.7% of Australian respondents (N=15) reported feeling a lack of opportunity in the department/institution due to their gender (P value = 0.039). Notably, overall 40% of female respondents (N=65) felt a lack of opportunity in their department/institution due to their gender.

### Discrimination

When asked about experiencing discrimination, 31% of non-white respondents (N=29) versus 10.8% of white respondents (N=83) reported experiencing discrimination within their lab and/or clinic (P value = 0.018). No significant difference for experiencing discrimination was found at the institutional level between non-white and white respondents, suggesting environments of discrimination experienced by non-white scientists may be most pronounced at the local lab/clinic level.

When questioned about witnessing discrimination in the lab/clinic, there was no statistically significant difference between ethnic/racial groups. 26.1% of respondents (N=111) witnessed discrimination in their local lab/clinic; 33.3% of white respondents (N=81) and 35.7% of non-white respondents (N=28) reported witnessing discrimination at their department/institution (**Table 1**).

In regard to gender, 36.4% of female respondents (N=66) versus 18.4% of male respondents (N=49) reported *feeling* discriminated against at their respective department/institution (P value = 0.03). In concordance with these findings, 43.3% of female respondents (N=67) versus 19.6% of male respondents (N=46) reported *witnessing* discrimination in the department/institution (P value = 0.009). No statistically significant results regarding an association between gender and feelings of discrimination were reported in the lab/clinic.

In regards to continent of origin, 37.1% of North American (N=70) and 57.1% of Australian respondents (N=14) reported witnessing discrimination in their department/institution, while only 5.3% of European respondents (N=19) shared these experiences (P value = 0.005) (**Table 1**). When categorizing responses to discrimination based on rank/position, the data indicates that 60% of junior faculty (N=20) reported feeling discriminated against in the department/institution in comparison to 26.9% of senior faculty/leadership (N=52) (P value = 0.003). Among all participants surveyed, 28.9% reported feeling discriminated against in their department/institution.

Overall, there was an elevated number of females in junior faculty than senior faculty positions among the respondents (not statistically significant). Among both senior and junior faculty, the percentage of females who reported feeling discriminated is about twice as much as males. When analyzed for each gender, the percentage who feel discriminated is twice as high in junior ranks compared to senior ranks (i.e., twice as many junior males feel discriminated vs senior males, twice as many junior females feel discriminated relative to senior females). Regardless of gender, junior faculty have a high proportion of feeling discriminated.

### Sexual Harassment

The topic of sexual harassment in academia has been recently pushed into the spotlight, with institutions taking action against individuals who abuse their power/position. We asked scientists in the melanoma community whether they experienced and/or witnessed sexual harassment in their lab/clinic or at their institution. Notably, 13.4% of female respondents (N=67), relative to no male respondents (N=48), reported *experiencing* sexual harassment in the department/institution (P value = 0.01) (**Table 1**). 12.9% and 27.8% of respondents have *witnessed* sexual harassment occurring in their lab/clinic (N=116) or department/institute (N=115), respectively.

If we look at this regarding gender, 30.3% of women (N=66) reported witnessing sexual harassment in the department/institution.

### Fairness in Promotions, Funding, and Raises

A total of 51% of respondents (N=98) reported feeling that promotions, funding, raises, and other opportunities are unfair in *their institution*, with 20 individuals not responding or preferring not to answer (**Table 1**). A large proportion of those surveyed believe that promotions, funding, raises, and other opportunities are not awarded fairly across the scientific community. Specifically according to gender, 89.1% of female (N=64) versus 72.1% of male respondents (N=43) reported feeling these notions (P value = 0.04); 82.2% of total respondents (N=107) expressed concerns of unfair promotions, funding, raises, and other opportunities across the scientific community. This sentiment appeared to fluctuate across continent of origin; 89.4% of North American (N=66), 85.7% of Australian (N=14), 61.1% of European respondents (N=18), and 1 of 3 respondents from other origins reported feeling promotions, funding, raises, and other opportunities were awarded unfairly across the scientific community (P value = 0.006) (**Table 1**).

It is worth noting that 17% of those surveyed preferred not to respond to the survey question regarding fairness in their institution. 54.2% of North American (N=59) and 50% of European respondents (N=16) reported feeling that promotions, funding, raises, and other opportunities are unfair in *their institution*.

## Discussion and Next Steps

Through our inaugural DEI Climate Survey, we have now begun to face the remnants of structural racism, inequity, and lack of inclusivity that persist in our melanoma community. As a community, we need to decide on best practices to address these concerns to protect the right to a diverse, equitable, and inclusive community each member of our community deserves. Our findings indicate that there are statistically significant differences in feelings and experiences regarding DEI amongst the SMR community. These results further reinforce the need for us to address ongoing oppression, bias, and exclusion faced by many SMR community members. Of note, the sample sizes relating to sexual orientation were too small to be representative of the SMR community, so the results were not included in the analysis. We hope with more survey responders in the next iteration that we would be able to more precisely dissect associations with discrimination, sexual harassment, and opportunity inequity. Overall, there appears to be an agreement among members of the SMR community regarding concerns to fair access and opportunity based on gender, ethnic/racial group, and geographic location.

